# Neural responses to sensory novelty with and without conscious access

**DOI:** 10.1101/2020.10.31.363366

**Authors:** Sergio Osorio, Martín Irani, Javiera Herrada, Francisco Aboitiz

**Affiliations:** Laboratorio de Neurociencia Cognitiva y Evolutiva, Departamento de Psiquiatría, Escuela de Medicina, Pontificia Universidad Católica de Chile, Santiago de Chile; Laboratorio Neurodinámicas de la Cognición, Departamento de Psiquiatría, Escuela de Medicina, Pontificia Universidad Católica de Chile, Santiago de Chile; Centro Interdisciplinario de Neurociencias, Pontificia Universidad Católica de Chile, Santiago de Chile; Instituto de Neurociencia Biomédica, Facultad de Medicina, Universidad de Chile, Santiago De Chile

**Keywords:** Conscious access, novelty, uncertainty, theta oscillations, P3, pupil response

## Abstract

Detection of novel stimuli that violate statistical regularities in the sensory scene is of paramount importance for the survival of biological organisms. Event-related potentials, phasic increases in pupil size, and evoked changes in oscillatory power in the theta (4-8 Hz) frequency range have been proposed as markers of sensory novelty detection. However, how conscious access to novelty modulates these different brain responses is not well understood. Here, we studied the neural responses to sensory novelty in the auditory modality with and without conscious access. We identified individual thresholds for conscious auditory discrimination and presented to our participants sequences of tones, where the last stimulus could be another standard, a subthreshold target or a suprathreshold target. Participants were instructed to report whether the last tone of each sequence was the same or different from those preceding it. Results indicate that stimulus evaluation and overt decision-making mechanisms, indexed by the P3 event-related response and reaction times, best predict whether a novel stimulus will be consciously accessed or not. In contrast, increased pupil size does not predict conscious access to novelty but reflects unexpected sensory uncertainty. These results highlight the interplay between bottom-up and top-down mechanisms and how the brain weights neural responses to novelty and uncertainty during perception and goal-directed behavior.

## Introduction

The ability to extract statistical regularities from sensory scenes and detect novel events that do not match self-generated predictions is of paramount importance for biological organisms. Novelty detection can simultaneously engage multiple bottom-up and top-down mechanisms that mediate autonomic and goal-directed responses which are crucial for survival (Anderson et al., 2012; Escera & Malmierca, 2014; Itti & Baldi, 2005; Ranganath & Rainer, 2003; Tiitinen et al., 1994). How the brain weighs such interplay between perception and cognition is in part determined by stimulus properties and brain states (Alamia et al., 2019; Näätänen et al., 2007; Polich, 1987; Teixeira et al., 2014). However, how conscious access (i.e., the availability of a reportable sensory experience for reasoning and rational decision making, Mashour et al., 2020; Naccache, 2018) modulates the neural responses triggered by the detection of sensory novelty is less understood.

The brain response to novelty in the auditory modality has been previously characterized by the sequence of two Event-Related Potentials (ERPs): the Mismatch Negativity (MMN) and the P3 response. Auditory stimuli that violate the predictions of the central auditory system elicit an MMN response at around 200 ms after odd stimulus onset. This event is independent of attentional states, and presumably reflects the automatic generation of sensory memory traces and detection of mismatching representations in early auditory cortices (Bekinschtein et al., 2009; Fischer et al., 1999; Garrido et al., 2009; Näätänen et al., 2007, 2011; Tiitinen et al., 1994). Novel auditory events that are attended to and consciously accessed additionally elicit a P3 response, an event that has been proposed to reflect attentional resource allocation, revision of stimulus-driven mental representations and memory-dependent processing leading to decision-making (Comerchero & Polich, 1999; Polich, 1987, 2007; Sutton et al., 1965). This response has been further dissected into two separate subcomponents that might reflect different stages of information processing. The P3a event, or novelty P3, can be observed in fronto-central electrodes at an early latency (250-300 ms) when subjects detect novel but task irrelevant stimuli. In contrast, the P3b event is observed at a later latency in centro-parietal electrodes when task-relevant novel stimuli are consciously reported. This has led to the proposal that the P3a reflects orienting attention and stimulus-evaluation processes, recruiting anterior cingulate and prefrontal cortical generators, while the P3b reflects subsequent information maintenance and deployment of working memory, recruiting parietal, postcentral and posterior cingulate generators (Comerchero & Polich, 1999; Polich, 2007; Ranganath & Rainer, 2003).

Two other potential markers of auditory novelty detection are the pupil dilation response and evoked oscillations within the theta (4-8 Hz) frequency range. Phasic increases in pupil size have been associated with the detection of auditory novelty, but whether this response reflects conscious access is contentious. Some studies have reported phasic increases in pupil size for auditory stimuli that are consciously reported but not for stimuli that escape conscious detection (Bala et al., 2020; Quirins et al., 2018). Others have shown that during passive listening conditions, only salient changes in the auditory scene elicit a pupil response. In contrast, when behaviorally relevant, any novel stimulus elicits a pupillary response (Liao et al., 2016; Zhao et al., 2019). This suggests that the pupil is highly sensitive to the behavioral demands imposed by the task at hand, and casts doubts on the idea that the pupil is a reliable window to the contents of consciousness. Additionally, previous magnetoencephalography (MEG) and electroencephalography (EEG) studies on auditory mismatch detection have reported increases in the power of theta oscillations concomitant with the MMN and the P3 at the scalp level, and in source-reconstructed prefrontal and temporal regions (Hsiao et al., 2009; Javitt et al., 2018; Recasens et al., 2018; Solís-Vivanco et al., 2021). However, none of these studies explicitly investigated how conscious access modulates oscillatory dynamics in response to novel auditory events.

Here, we investigated how conscious access modulates the neural responses to sensory novelty in the auditory modality. We identified individual thresholds for conscious discrimination and presented to our participants sequences of standard tones of variable length. Participants were asked to decide whether the last stimulus was the same or different from previous stimuli in the sequence. The last tone could be another standard tone (tgtSTD), a subthreshold deviant target (subDEV), or a suprathreshold deviant target (supraDEV). We resorted to signal detection theory to study stimulus detectability and response systematicity to the different types of targets. We then aimed to replicate well-established findings in ERP literature about the MMN and P3 as reliable markers of conscious access to sensory novelty. Next, we investigated how evoked pupil responses and changes in theta power are modulated by conscious access to auditory novelty. Finally, we examined which of these behavioral, electrophysiological and pupillometric variables best predicted whether novel auditory stimuli would be consciously accessed.

## Methods

### Participants

Twenty-eight right-handed subjects (16 females) with no self-reported record of auditory, neurological, or neuropsychiatric disorders voluntarily agreed to participate in this study (mean age = 25.92, SD = 3.10). All participants reported normal hearing and normal or corrected-to-normal vision. Subjects with extensive and/or formal musical training (> year) were not considered for the study. Participants were recruited from among the undergraduate and postgraduate community at Pontificia Universidad Católica de Chile and Universidad de Chile. Protocol and procedures were reviewed, approved, and supervised by the ethics committee for life and medical sciences at Pontificia Universidad Católica de Chile.

### Procedures and stimuli

Participants sat 50 centimeters away from of a screen within a Faraday cage in a silent, dimly-lit room. Brain activity was recorded using a 64-channel Biosemi EEG system and pupil size was recorded using an Eyelink 1000 eye-tracker calibrated at the beginning of the experimental session. 150-millisecond long narrowband sinusoidal tones were presented binaurally via special airtube earphones (ER-1 Etymotic Research) with an interstimulus interval of 150 ms. All stimuli were created using Audacity and set to be delivered at an intensity of 70dBs. The task was programmed using Presentation (NeuroBehavioral Systems).

The experiment included a 15-minute training session and three experimental blocks of approximately 15 minutes each. At the beginning of each experimental block, participants performed a staircase procedure (supplementary materials, figure s1). This procedure allowed identifying subject-specific conscious discrimination thresholds and setting subthreshold tonal stimuli adaptively and according to individual hearing abilities. For the staircase procedure, a set of sixteen tonal stimuli were used per block with a base frequency of 800Hz, 1000Hz or 1200Hz (supplementary materials, table s2). Then, participants would perform one of the three blocks of the main task. Each block in the main task included 270 trials, for 90 targets per condition. Nine participants performed a longer version of the task (320 trials, 120 targets per condition).

In the main task (figure 1), participants were instructed to listen carefully to a series of standard tones and decide whether the last stimulus in each sequence was the same or different from the preceding stimuli. The number of standard tones before each target randomly varied between three and five tones. Participants reported their decision by pressing one of two buttons upon appearance of a prompt on screen 1,000 milliseconds after target onset. There was no time limit for response and participants were told to prioritize response accuracy over response speed. For each block, standard tones were presented at one of the base frequencies used for the staircase procedure. Target stimuli could be either another standard tone (tgtSTD), a tone that was 50 Hz above the base frequency (supraDEV) or a tone that was below each participants’ discriminatory threshold (subDEV) identified during the staircase procedure before the beginning of the block. The theoretical probability for each type of target was 33.333%.

**Figure 1.**
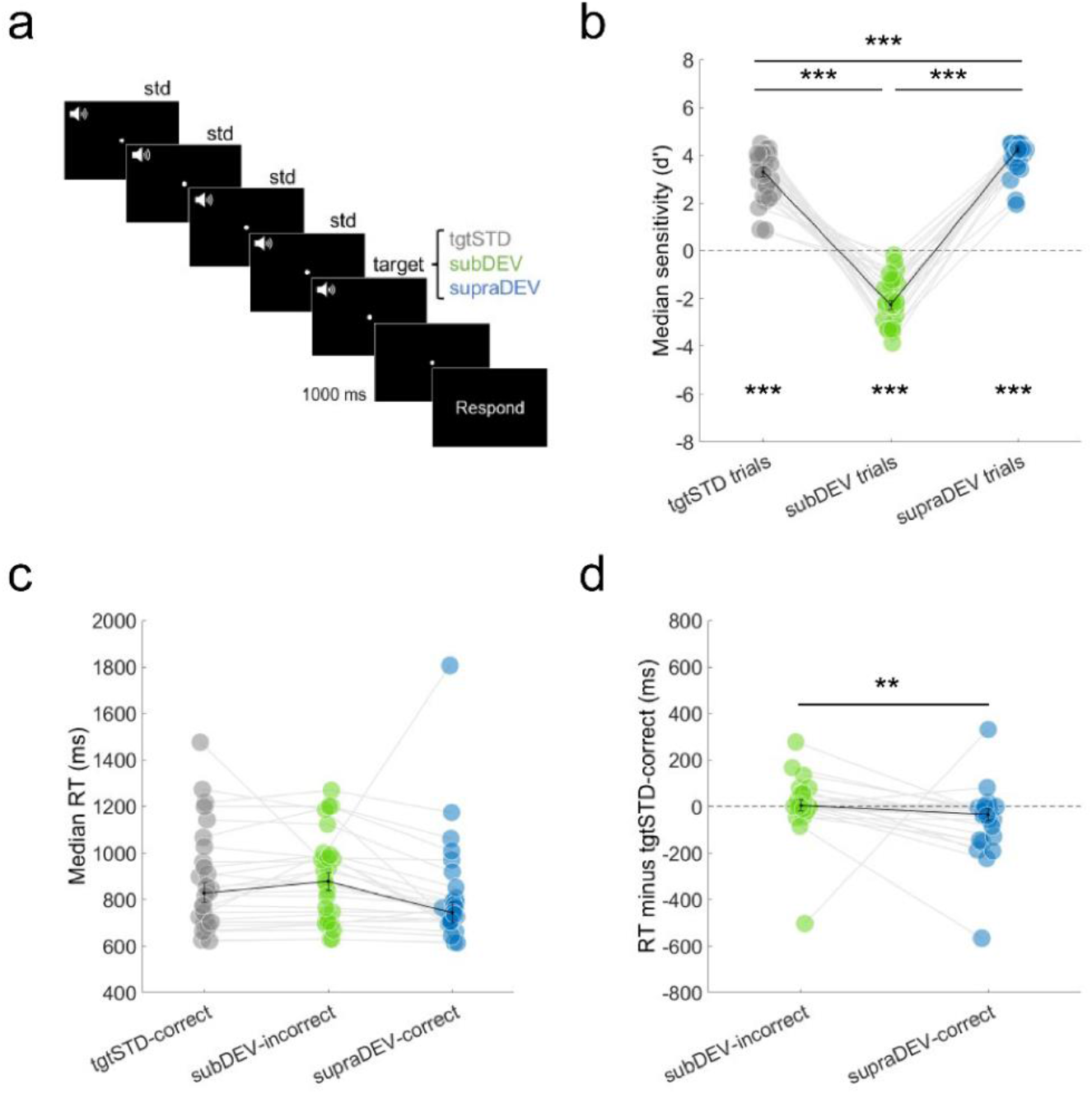
Main task and behavioral results. **a**. Schematic representation of a single trial in the main task. Participants had to decide whether the last of a sequence of tones was the same or different from the preceding tones. The target could be another standard, a subthreshold or a suprathreshold stimulus. The number of standard tones preceding the target randomly varied between three and five. Participants responded by pressing one of two buttons when a prompt appeared on screen, 1,000 ms after target onset. **b**. Sensitivity values for the three different types of targets. Whiskers represent the median and the standard error (S.E.) of the median. Asterisks at the bottom represent a significant difference from zero. Asterisks with bars represent a significant difference across conditions (*** < 0.001, ** < 0.01). **c**. Median reaction times for tgtSTD-correct, subDEV-incorrect, and supraDEV-correct trials. **d**. Median RT difference for subDEV-incorrect and supraDEV-correct compared to tgtSTD-correct trials.

### Behavioral data analysis

Behavioral data was obtained using Presentation (NeuroBehavioral Systems, NBS). Default logfiles were preprocessed and analyzed using in-house MATLAB scripts. We resorted to Signal Detection Theory (Green & Swets, 1966) to estimate sensitivity indices (*d’*). Sensitivity is a measure of target detectability (signal) that considers both error rates (noise) and response systematicity (bias), thus outperforming raw accuracy and hit probabilities in accounting for human decision-making processes in perceptual tasks (Kingdom & Prins, 2016). Sensitivity (*d’*) values were calculated for each target type by measuring the distances between mean number of correct responses and incorrect responses in standard deviation units, while correcting for perfect rates according to the 1/2N rule (Green & Swets, 1966; Hautus et al., 2022). Sensitivity values were obtained per block and then averaged across blocks per participant. Positive *d’* values suggest a higher proportion of correct over incorrect responses, whereas negative values suggest the opposite pattern. The farther away from zero, the lesser the probability that performance is due to chance. Data from one participant was rejected from any further analysis due to an erratic performance during the staircase procedure which resulted in unreliable discriminatory thresholds. Except for sensitivity, all subsequent analyses were carried out for tgtSTD and supraDEV trials followed by correct responses, and for subDEV trials followed by incorrect responses (percent rejected: tgtSTD, mean = 2.04, SD = 10.24; subDEV, mean = 16.76, SD = 29.31). Reaction times were calculated by subtracting the 1,000 ms time-window between target onset and response prompt from the participant’s actual response time. Any negative reaction time was considered a false alarm and was therefore rejected, along with outliers below and above the 2.5 and 97.5 percentiles in the data (percent rejected: tgtSTD-correct, mean = 3.70, SD = 0.96; subDEV-incorrect, mean = 3.02, SD = 1.22; supraDEV-correct, mean = 3.82, SD = 0.77).

### EEG data preprocessing

Electroencephalographic data was acquired at a sampling rate of 1024 Hz. After acquisition, the signal was preprocessed using Brainstorm (Tadel et al., 2011). The continuous EEG signal was linearly detrended and noisy segments were rejected. Next, the EEG signal was re-referenced to the average of all electrodes and band-passed filtered between 1 and 45Hz using an FIR Keiser filter (Filter order = 37124, Stopband attenuation = 60dB) as implemented in the Brainstorm toolbox. Oculomotor and blink-related artifacts were removed using and Independent Component Analysis (Makeig et al., 1996) on the continuous EEG signal (mean components rejected = 8.77, SD = 2.69). Data was subsequently epoqued between -2500ms and after 2500ms upon target stimulus presentation. A trial-by-trial inspection was carried out to visually identify and reject bad trials and channels (mean = 1.00, SD = 1.15). Next, an automatic trail rejection procedure was performed, marking as bad any trial where EEG signal exceeded 100 microvolts in amplitude (mean = 7.25, SD = 7.15 including manual and automatic rejection). Mean Event Related Potentials were computed as the baseline-corrected average across subjects for tgtSTD-correct, subDEV-incorrect, and supraDEV-correct trials. Baseline correction was applied by subtracting the mean ERP between -500 milliseconds and time zero. Time-frequency decomposition analyses were conducted using the fieldtrip toolbox (Oostenveld et al., 2011) by means of a 7-cycle wavelet in steps of 0.5 Hz between 1 and 30 Hz, for a time window of -1500 to 1500. The grand average for tgtSTD-correct, subDEV-incorrect, supraDEV-correct trials was obtained and dB-normalized using a baseline period of -500 to -100 ms. EEG data from one participant was incomplete due to technical problems during acquisition and had to be rejected from analyses.

### Pupillometry

Pupil data was acquired using Eyelinks’ acquisition software at a sampling rate of 1,000 Hz. Calibration procedures were carried out at the beginning of the experiment. Pupil area, horizontal and vertical gaze positions were recorded from the right eye of each participant. Blinks and gaze artifacts were detected by Eyelinks’ default algorithms. Pupil data was preprocessed using Anne Urai’s pupil toolbox (Urai et al., 2017) plus additional custom MATLAB scrips. Eyelink-defined and additionally detected blinks were padded by 150 milliseconds and linearly interpolated. The pupil response evoked by blinks and saccadic events was identified via deconvolution and removed using linear regression (Knapen et al., 2016). The signal was then filtered between 0.01 Hz and 10 Hz using a second-order Butterworth filter and then down sampled to 250 Hz. Data was epoqued between 2500 milliseconds before and 2500 milliseconds after the onset of target stimuli. Data was subsequently z-normalized and the grand average of the pupil size for tgtSTD-correct, subDEV-incorrect, and supraDEV-correct trials was estimated. The time window for baseline correction comprised -500 milliseconds to time zero. Pupil data from three participants was unavailable due to problems during data acquisition.

### Statistics

Non-parametric permutation statistics for behavioral data were implemented using custom-made MATLAB scripts. For *d’* and reaction times, we used a one-sample t-tests and non-parametric permutation procedures. For permutation procedures, the median values across conditions were obtained from the observed data. Next, we created a null distribution by exchanging condition labels over each iteration of a 10,000-permutation procedure. Then, observed values were compared against the percentiles corresponding to the Bonferroni-corrected alpha values obtained from the null distribution, and corresponding p-values were calculated. Bootstrapping procedures were similarly implemented, except that instead of interchanging labels across conditions, n values were randomly sampled (with replacement) from the observed values per condition for each iteration. For time series data such as EEG and Pupil ERPs, we implemented the same procedure over time; that is, sample-by-sample with an additional criterion that for an effect to be significant, below-threshold p values should be observed for at least three continuous samples. For event-related potentials, we performed this analysis within *a priori* selected time windows, corresponding to the approximate latency of the MMN (150-250 ms) and the P3 (280-380 ms). Given our strongly directional hypotheses, we used one-tailed testing and an alpha level of 0.01. For pupil data, however, we did not have a strong directionally hypothesis or predefined time windows of interest informed by the literature, and therefore performed two-tailed analyses between 0 and 1500 ms. For time-frequency data, two-tailed cluster-corrected statistics were obtained via a non-parametric, dependent samples permutation procedure (10,000 permutations) using the Fieldtrip toolbox (Oostenveld et al., 2011) between 150-250 ms and 280-380 ms. For all cluster-statistics, clustering of neighboring channels was set using the triangulation method and the uncorrected p-value for cluster inclusion was set at p < 0.05. Cluster significance probability was corrected using the Monte-Carlo method (α = 0.05), for 10,000 permutations and a minimum number of neighboring channels of three. All Spearman correlation analyses were implemented using MATLAB’s statistics toolbox default function (two-tailed testing).

Multiple logistic regression analysis was implemented using R 4.1.3. (R Core Team, 2021). We extracted the individual and mean or median values for our predictors, including mean *d’*, mean *d’* difference, median RT, median RT difference, mean pupil size, mean pupil size difference, mean ERP between 150-250 and 280-380 ms (for channel Cz), mean theta power between 150-250 and 280-380ms, and mean theta power difference between 150-250 and 280 ms (averaged across electrodes in statistically significant clusters). Subjects with missing data were rejected from this analysis. An initial model was created using the maximum number of predictors that would not result in model overfitting. We next created another model by rejecting the predictor with the highest p value in the summary statistics report. In some instances, we replaced a predictor by its mean difference to test how this affected the initial model and to confirm our decision to leave this predictor out in the subsequent models. We followed this procedure until we reached single-predictor models. For each model (n = 15), we estimated the Akaike Information Criterion (AIC) and the Bayesian Information Criterion (BIC). Lower AIC and BIC values are indicative of a better fitting model (supplementary materials, table s2). This resulted in one model showing the lowest AIC and BIC values (model 7).

## Results

### Behavioral results

We estimated sensitivity indices (*d’*) for each type of target across blocks and participants. We expected systematic and correct identification of non-novel standards (tgtSTD) and suprathreshold deviant targets (supraDEV), which should be reflected by positive *d’* values. In contrast, we expected participants to systematically and incorrectly judge subthreshold deviant targets (subDEV) as standard tones, which should be reflected by negative *d’* values. Figure 1b shows individual and group median sensitivity values for the three types of target stimuli. In line with our expectations, tgtSTD (*d’* = 3.30, SD = 0.99) and supraDEV targets (*d’* = 4.26, SD = 0.66) were associated with positive *d’* values, whereas subDEV targets were associated with negative values (*d’* = -2.27, SD = 0.96). A one sample t-test (n = 27, two-tailed) showed that median sensitivity values statistically differed from zero (tgtSTD, t = 16.48, p < 0.001; subDEV, t = 31.05, p < 0.001; supraDEV, t = -11.31, p < 0.001). This suggests that, on average, supraDEV targets were consciously accessed as novel, and that subDEV targets were subconsciously processed, but not consciously accessed as novel. A non-parametric permutation procedure showed that the median *d’* statistically differed across conditions (figure 1b, n = 27, 10,000 permutations, two-tailed, Bonferroni-corrected, p > 0.001).

We next investigated reaction times (RT) to trials conforming to these target-response combinations. Figure 1c shows the median RT to tgtSTD-correct (median = 828.30, SD = 222.65), subDEV-incorrect (877.30, SD = 189.58) and supraDEV-correct trials (median = 742.90, SD =238.57). We then subtracted the median RT to tgtSTD-correct trials from the other two conditions. This allowed us to understand whether decision-making processes to subDEV-incorrect and supraDEV-correct trials were slower or faster compared to a baseline condition. Results showed that RTs to subDEV-incorrect trials were marginally slower (4.5 ms) than the median RTs to tgtSTD-correct, whereas supraDEV-correct targets were, on average, 37.45 ms faster than the median RT to tgtSTD-correct trials. A permutation test showed that the median RT difference for subDEV-incorrect and supraDEV-correct trials significantly differed at the group level (n = 27, 10,000 permutations, two-tailed, p = 0.0016, figure 1d).

### EEG results

We then computed the mean event-related responses (ERPs) to tgtSTD-correct, subDEV-incorrect, and supraDEV-correct trials (figure 2). We focused our analyses in two predefined time windows: 150-250 and 280-380 ms, which coincide with the approximate latency of the MMN and the P3 events. We expected to observe a MMN response to both subDEV-incorrect and supraDEV-correct trials, and a P3 response only to supraDEV-correct trials. As hypothesized, a bootstrapping procedure (n = 26, one-tailed, 10,000 bootstraps, p < 0.01) showed that the mean ERP response recorded at electrode Cz in response to subDEV-incorrect and supraDEV-correct targets was more negative compared to tgtSTD-correct trials (figure 2a). This difference was statistically significant between 172 and 200 ms for subDEV-incorrect targets and between 152 and 192 ms for supraDEV-correct targets. In contrast, only supraDEV-correct targets elicited a positive deflection in the ERP wave between 280 and 380 ms. Similar results were observed in other centro-parietal electrodes, including C2 and CPz (figure 2b and 2c), with the amplitude of the P3 decreasing from anterior to posterior. At 180 ms, ERP topographies showed bilateral temporal positivities and frontal negativities consistent with scalp ERP activation associated with the MMN response (figure 2d). In turn, centro-frontal positivities at 320 ms were also consistent with the P3 event. The observed fronto-central electrode distribution of this neural response is consistent with the topographical characteristic of the P3a event.

**Figure 2.**
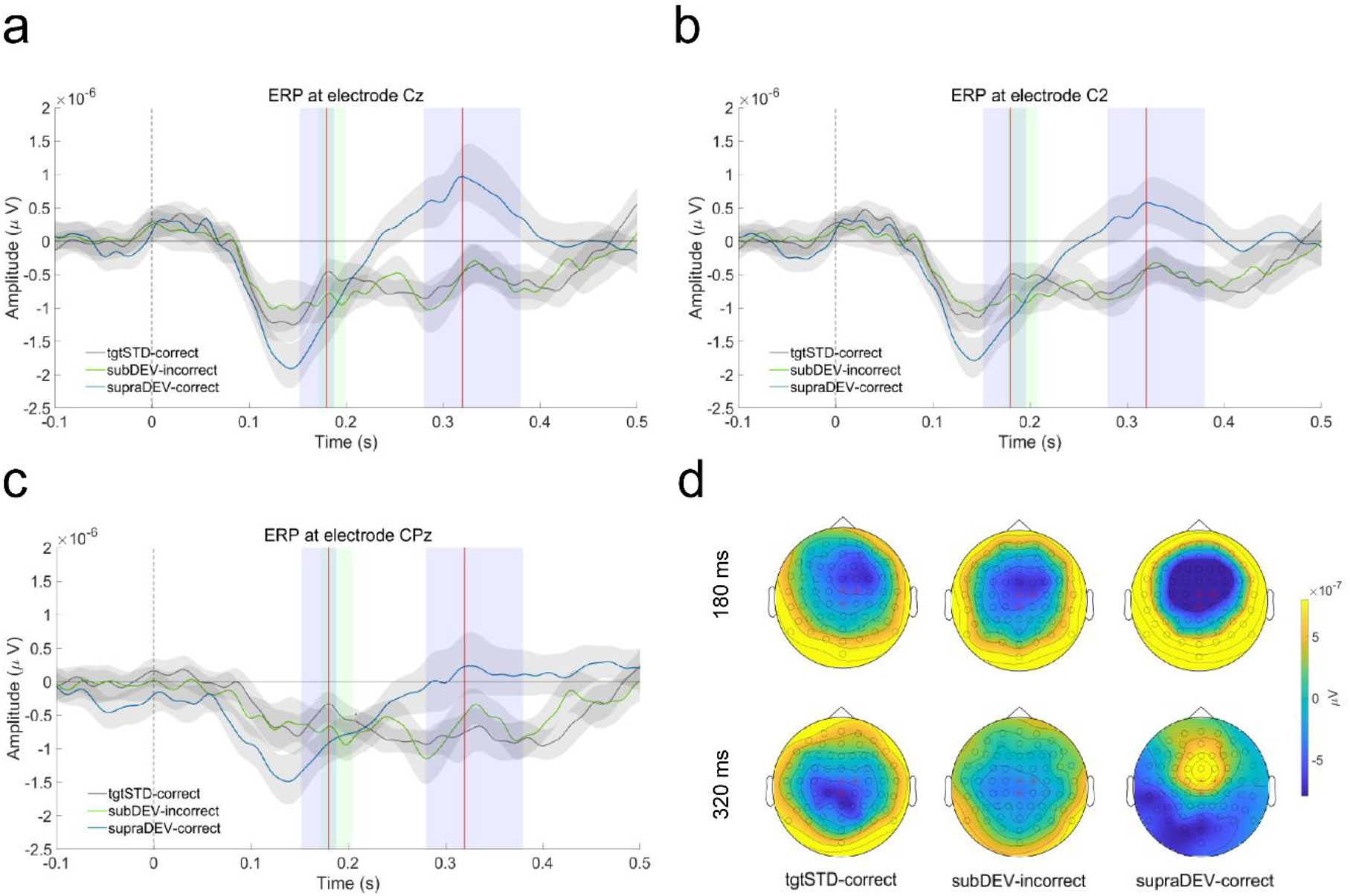
Event-related potentials (ERPs). Mean ERP measured at electrodes Cz (**a**), C2 (**b**) and CPz (**c**). Gray shades represent the 95% C.I. of the mean. Colored shades represent a statistically significant difference between subDEV-incorrect (green) and supraDEV-correct (blue) trials against tgtSTD-correct trials at an alpha level of 0.01. **d**. Topographical maps showing ERP activation at 180 ms (top) and 320 ms (bottom), also indicated by red lines in the time-series data. Red asterisks represent the location of electrodes plotted in a, b, and c.

### Pupillometry

We later investigated the evoked pupil responses to tgtSTD-correct, subDEV-incorrect, and supraDEV-correct trials. If the pupil response reflects conscious access to auditory novelty, we should observe a pupil response to supraDEV-correct, but not to subDEV-incorrect trials. In sharp contrast with this expectation, results (n = 23) showed that all types of targets (including tgtSTD targets) elicited a phasic increased in pupil size, peaking at around 1,200 m after stimulus onset regardless of whether they had been consciously accessed or not (figure 3a). After subtracting the mean pupil response to tgtSTD-correct trials from the other two conditions, pupil size for subDEV-incorrect was significantly higher compared to supraDEV-correct trials between 164 ms and 512 ms (10,000 bootstraps, two-tailed, p < 0.016, figure 3b). We also studied the relationship between pupil size difference within this time window and the mean ERP values measured at electrode Cz. For supraDEV-correct trials, increased pupil size was associated with lower amplitude of the MMN and higher amplitude of the P3 supraDEV-correct trials within 150-250 ms (n = 22, rho = 0.43, p = 0.048, figure 3c, right) and 280-380 ms (n = 22, rho = 0.47, p = 0.027, figure 3d, right).

**Figure 3.**
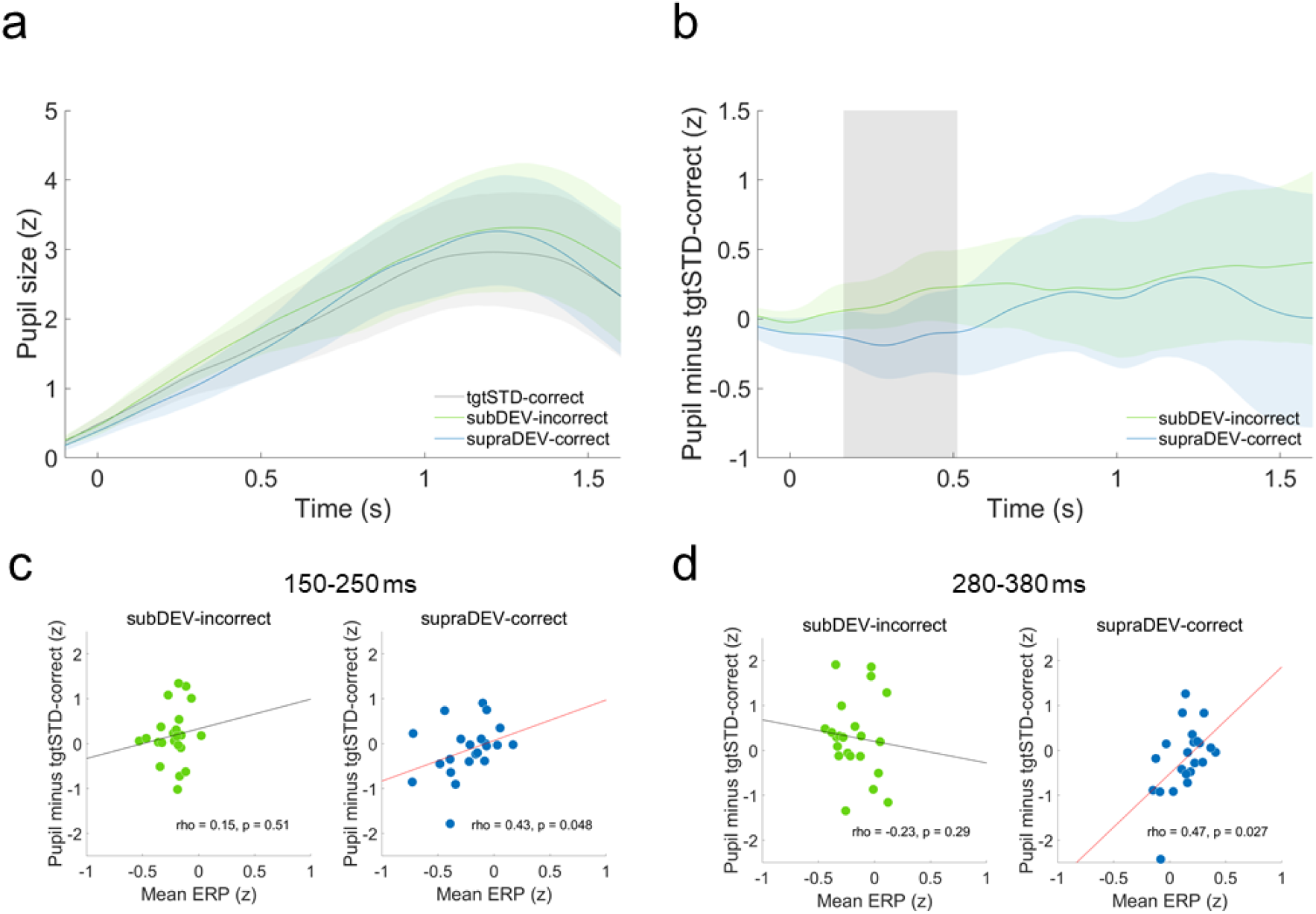
Pupillometry. **a**. Z-normalized mean pupil response for the three types of trials. Colored shades represent the 95% Confidence Intervals (C.I.) of the mean. **b**. Mean pupil difference between subDEV-incorrect and supraDEV-correct trials against tgtSTD-correct trials. Gray shade represents a statistically significant difference between conditions. **c**. and **d**. Spearman correlations between mean pupil difference against tgtSTD-correct trials and mean ERP response to subDEV-incorrect and supraDEV-correct trials between 150-250 ms (**c**) and between 280-380 ms (**d**). Red best fit lines illustrate a statistically significant correlation.

**Figure 3.**
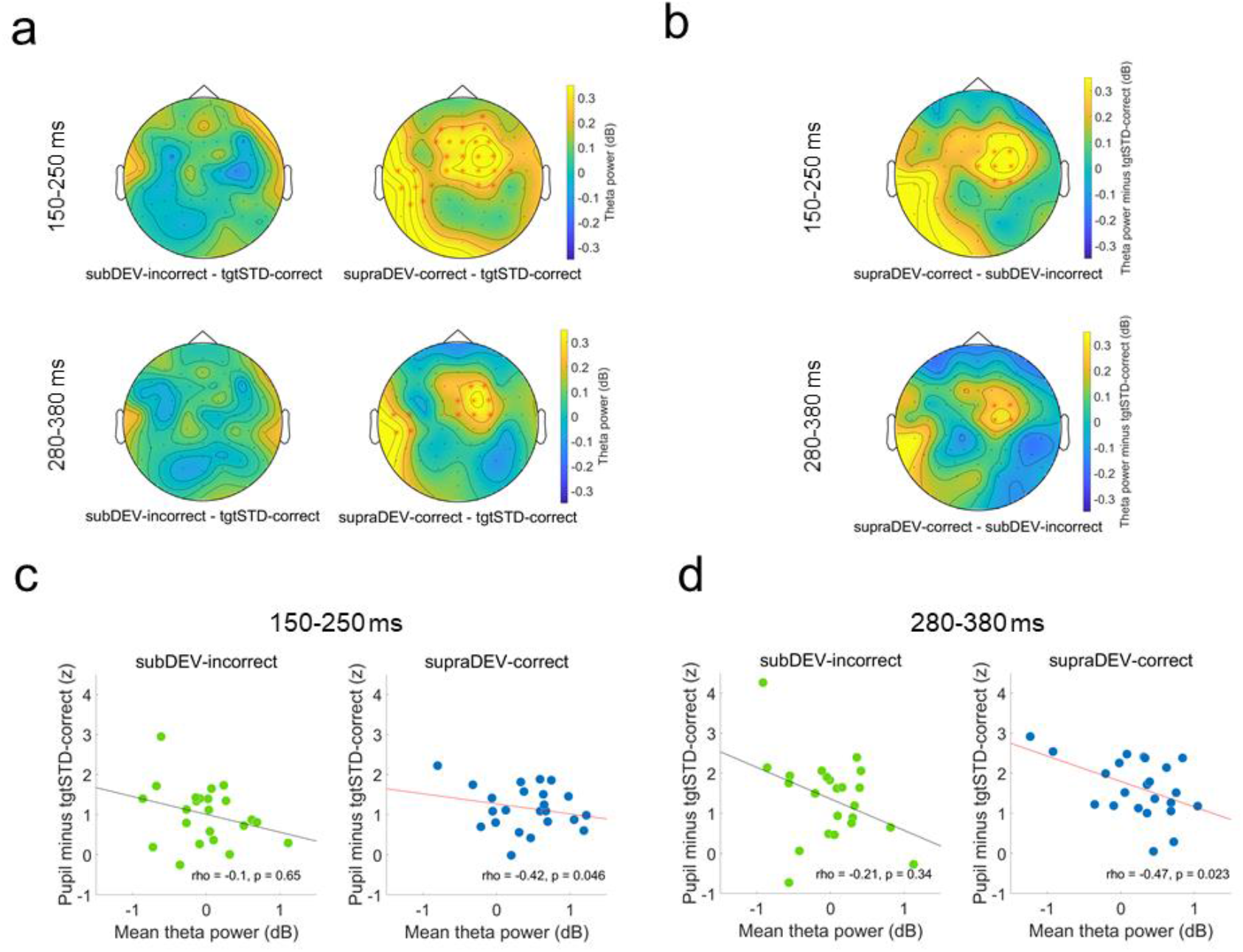
Wavelet time-frequency analysis. **a**. Mean power difference between subDEV-incorrect (left column) and supraDEV-correct (right column) against tgtSTD-correct trials between 150-250 ms and 280-380 ms. Electrodes in red represent clusters showing a statistically significant difference. **b**. Mean theta power difference between supraDEV-correct and subDEV-incorrect trial, after subtraction of tgtSTD-correct trials. **c**. and **d**. Spearman correlation between mean pupil difference against tgtSTD-correct and mean theta power. Red lines represent a statistically significant correlation.

### Time-frequency analyses

Next, we studied evoked changes in theta (4-8 Hz) power in response to the different types of trials (figure 3). If theta power reflects conscious access to novelty, we should observe a power increase in this frequency band for supraDEV-correct trials, but not for subDEV-incorrect trials. As hypothesized, a non-parametric permutation test (n = 26, 10,000 permutations, two-tailed, cluster-corrected) showed that supraDEV-correct trials were associated with significant increases in theta power compared to tgtSTD-correct trials between 150-250 ms for a positive cluster of centro-frontal and left temporal distribution (t = 129.36, p < 0.001), and between 280-380 ms for two positive clusters, the first of centro-frontal distribution (t = 13.382, p = 0.020) and the second of left temporal distribution (t = 4.67, p = 0.034, figure 3a). No similar effect was observed for subDEV-incorrect trials compared to tgtSTD-correct trials. We also investigated any difference between supraDEV-correct and subDEV-incorrect trials after subtracting RT to tgtSTD-correct trials. Results showed that this difference was restricted to a cluster of electrodes with centro-frontal distribution, both between 150-250 (t = 42.58, p = 0.004) and between 280-380 ms (t = 4.81, p = 0.038, figure 3c). We therefore estimated the average power across the electrodes contained in the biggest cluster and investigated its relationship with pupil size difference. For supraDEV-correct trials, spearman correlations (n = 22, two-tailed) showed that increased theta power is associated to decreased pupil size. This relationship was statistically significant between 150-250 ms (rho = -0.42, p = 0.046) and between 280 and 380 ms (rho = -0.47, p = 0.023).

### Multiple logistic regression

As a final step, we investigated which of our variables of interest best predicted whether novel auditory stimuli would be consciously accessed. For this, we implemented a binomial multiple regression analysis using our behavioral, pupillometric and electrophysiological variables as predictors. We created a set of fifteen logistic models with varying numbers of predictors and used the Akaike Information Criterion (AIC) and the Bayesian Information Criterion (BIC) to find the best fitting model within this set (see methods and supplemental information, table s2). A model including RT difference, mean pupil size and mean ERP values between 280-380 ms best predicted conscious access to auditory novelty (AIC = 25.752, BIC = 31.863, McFadden’s pseudo *R*^2^ = 0.725, p < 0.001, table 1).

**Table 1.**
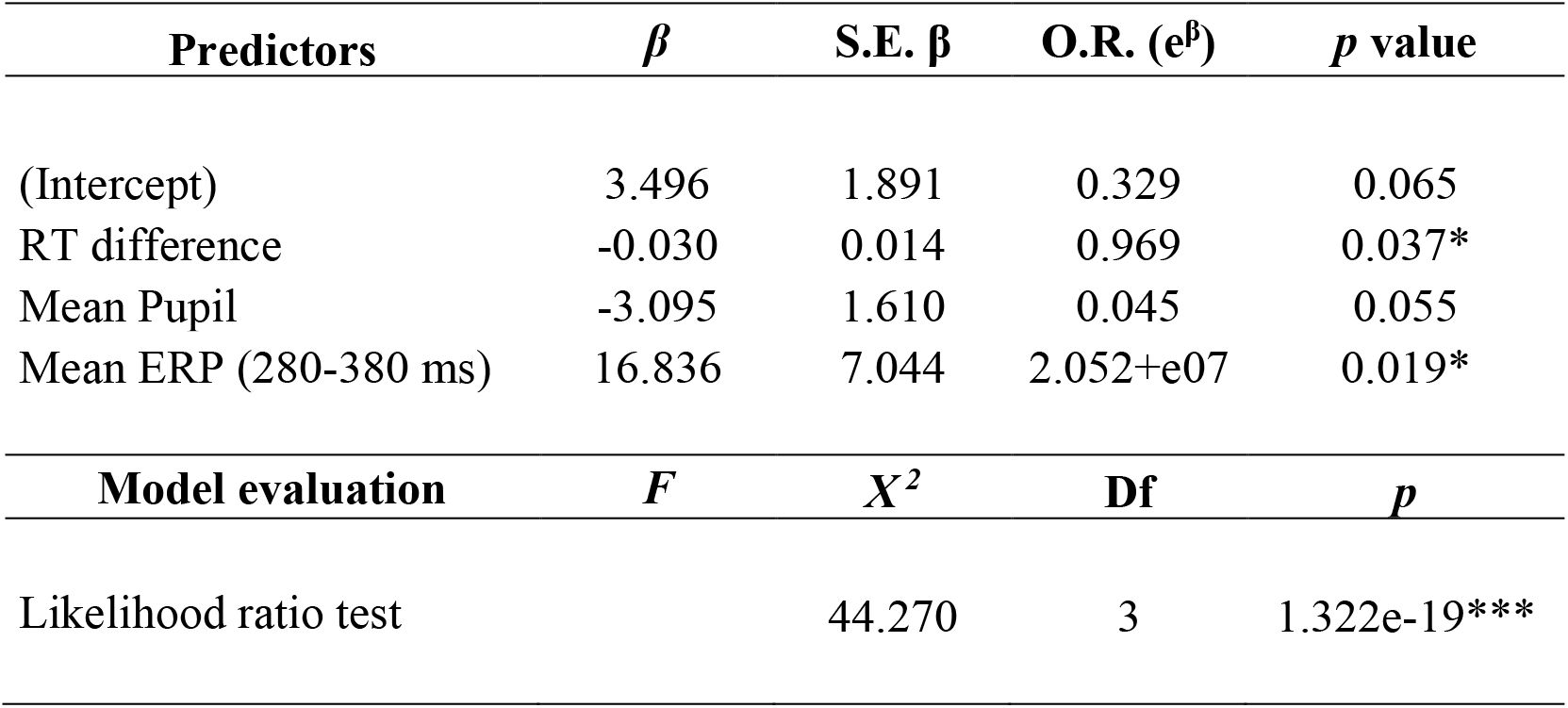
Best fitting logistic model predicting conscious access to auditory novelty and model evaluation against a null model. Asterisks indicate statistical significance at an alpha level of 0.05 (*), 0.01 (**) and 0.001 (***).

| Predictors | $\beta$ | S.E. $\beta$ | O.R. ($e^\beta$) | $p$ value |
| --- | --- | --- | --- | --- |
| (Intercept) | 3.496 | 1.891 | 0.329 | 0.065 |
| RT difference | -0.030 | 0.014 | 0.969 | 0.037* |
| Mean Pupil | -3.095 | 1.610 | 0.045 | 0.055 |
| Mean ERP (280-380 ms) | 16.836 | 7.044 | 2.052+e07 | 0.019* |

| Model evaluation | $F$ | $X^2$ | Df | $p$ |
| --- | --- | --- | --- | --- |
| Likelihood ratio test |  | 44.270 | 3 | 1.322e-19*** |

Results indicate that the log of the odds for of a novel stimulus being consciously accessed is negatively related to median RT difference (*β* = -0.030, p = 0.037) and mean pupil size (*β* = -3.095, p = 0.055), and positively related to the mean ERP between 280 and 380 ms (*β* = 16.836, p = 0.019). This is to say, the faster the reaction time and the higher the amplitude of the P3 response, the more likely that novel auditory stimuli are consciously accessed. A likelihood ratio test (*Χ* ^2^ = 44.270, df = 3, p < 0.001) confirmed that this model provides a better fit to the data over a null (i.e., intercept only) model.

## Discussion

### Behavioral and electrophysiological validation of the experimental paradigm

In this study, we investigated the neural responses to sensory novelty in the auditory modality with and without conscious access. We presented sequences of tones, where the last stimulus could be another standard tone, a subthreshold target, or suprathreshold target. Participants had to decide whether targets were novel or non-novel. As expected, subDEV targets were systematically reported as standard tones, reflecting subconscious processing but not conscious access to novelty (figure 1b). Systematic and correct identification of supraDEV targets, instead, reflected conscious access to a novel auditory stimulus (figure 1b). By subtracting reaction time to non-novel targets, we found that subDEV stimuli were responded to more slowly and supraDEV targets were responded to faster compared to tgtSTD-correct trials (figure 1d). Thus, overt decision making was sped up when the novel stimulus was above participants’ thresholds and was delayed when the stimulus was below threshold. This is in line with previous evidence showing how stimulus saliency and uncertainty facilitate and hamper behavioral responses, correspondingly (Garcia et al., 2017; Kayser et al., 2005; Perugini et al., 2016; Teixeira et al., 2014).

Previous studies have shown that subconscious processing of auditory novelty is indexed by the MMN negativity, a neural event that reflects early sensory memory mechanisms (Bekinschtein et al., 2009; Fischer et al., 1999; Garrido et al., 2009; Näätänen et al., 2007, 2011; Tiitinen et al., 1994), and that conscious access to novelty is characterized by the P3, an event that reflects attentional allocation, stimulus evaluation, and memory-dependent decision-making processes (Comerchero & Polich, 1999; Nieuwenhuis et al., 2005; Polich, 1987, 2007; Ranganath & Rainer, 2003; Sutton et al., 1965). We successfully replicated these findings by showing that both subthreshold and suprathreshold novel stimuli elicited a MMN response between approximately 180-200 ms, regardless of whether they were consciously accessed as novel (figure 2a, 2b, 2c). However, only consciously accessed suprathreshold novel stimuli elicited a fronto-central positivity between 280 and 380 ms (figure 2a, 2b, 2c) that is consistent with the P3 response (Comerchero & Polich, 1999; Escera & Malmierca, 2014; Ranganath & Rainer, 2003). What is more, the topographical distribution of this response matches that of the P3a event. This reflects the engagement of evaluation of stimulus evaluation mechanisms by higher-order processors, presumably in prefrontal and anterior cingulate regions (Garrido et al., 2009; Näätänen et al., 2007). Strikingly, we did not observe a shift to parietal electrodes in later latencies, as would be expected from a P3b response. Since we introduced a forced response delay in our task (figure 1a), it could be that memory-dependent decision-making processes underlying the P3b jittered from trial to trial, which could have obliterated the parietal subcomponent of the P3 complex. In spite of this, our behavioral and electrophysiological data do suggest that, unlike subthreshold stimuli, suprathreshold targets triggered the cascade of neural events that characterizes conscious access to novelty (Dehaene et al., 2006; Mashour et al., 2020; Naccache, 2018). We therefore take this as evidence that our task successfully teased apart subconscious processing from conscious access to novel auditory events.

### Pupil responses reflect sensory uncertainty and reallocation of attention to behaviorally relevant stimuli

Previous findings suggest that the pupil response reflects conscious detection of auditory novelty and regularity violations (Bala et al., 2020; Quirins et al., 2018), whereas other studies suggest that it reflects task-specific behavioral demands (Liao et al., 2016; Zhao et al., 2019). Our results align with this second line of evidence: first, all targets elicited a phasic increased in pupil size peaking at around 1,200 ms after target onset, regardless of stimulus identity or conscious access (figure 3a). This confirms that, similar to the P3 response, the pupil signals the reallocation of attentional resources to a behaviorally relevant stimulus (Liao et al., 2016; Zhao et al., 2019). Additionally, when the mean pupil response to non-novel targets was subtracted from the other two conditions, we observed increased pupil size to subthreshold targets between approximately 164 and 512 ms (figure 3b). This implies that the pupil response is also sensitive to unexpected uncertainty, which is in agreement with previous studies in both visual and auditory modalities (Alamia et al., 2019; Lavín et al., 2014; Urai et al., 2017; Zhao et al., 2019). This would also support the view that the pupil reflects phasic NE-mediated adaptation of arousal levels during goal-oriented and adaptive behavior (Aston-Jones & Cohen, 2005; Cockburn et al., 2021; Vazey et al., 2018). In contrast, no evidence was found that the pupil response can be considered a reliable marker of conscious access to novelty. We therefore propose that pupillary responses reflect the contribution of two separate components: an early component reflecting phasic arousal adaptation driven by uncertainty in the sensory scene and a later component reflecting stimulus-driven attentional reallocation to behaviorally relevant sensory events. For consciously accessed novel stimuli, we also found a positive association between pupil size difference and the P3 response (figure 3d). This fits evidence that both P3 and pupil responses in part depend on NE modulation and reflect phasic changes in arousal levels (Aston-Jones & Cohen, 2005; Masson & Bidet-Caulet, 2019; Nieuwenhuis et al., 2005). Our results also show that an increased pupil size is associated with smaller amplitude of the MMN (figure 3c). This finding reflects the antagonistic nature of these two neural mechanisms: increased stimulus saliency enhances sensory memory traces and facilitates automatic error detection indexed by the MMN but reduces uncertainty in the sensory scene, which would be reflected by reduced arousal adaptation. Indeed, previous studies have shown that MMN amplitude scales with contrast-based saliency (Sams et al., 1985; Tiitinen et al., 1994) and pupil responses increase with higher uncertainty (Alamia et al., 2019; Lavín et al., 2014; Urai et al., 2017).

### Theta power reflects information maintenance during conscious access to novelty

Increases in theta power following the detection of novel auditory stimuli have been previously observed in latencies consistent with the MMN and the P3 (Hsiao et al., 2009; Javitt et al., 2018; Recasens et al., 2018; Solís-Vivanco et al., 2021). We replicated these findings by showing that detection of suprathreshold deviants is related to increased theta power between 150-250 and 280-380 ms (figure 4a). We also extend these findings by showing that such theta-band power increase occurs only when novelty is consciously accessed. This confirms the role of theta oscillations in supporting conscious perception (Haque et al., 2020; Klimesch et al., 2001; Slagter et al., 2009). For supraDEV-correct trial, increased pupil size was correlated with reduced theta power in both time windows of interest (figure 4c and 4d). Similar to the Pupil-MMN relationship, phasic changes in pupil size and evoked oscillatory power within the range of theta reflect opposing cognitive mechanisms, theta power presumably marking conscious access to novelty. However, logistic regression analyses show that theta power does not predict increased odds of a novel stimulus being consciously accessed (table 1). Hence, rather than conscious access per se, an alternative is that theta power reflects information maintenance about consciously-accessed novel stimuli which would be necessary for accurate decision-making.

### Stimulus evaluation and overt decision-making mechanisms predict conscious access to novelty

Our logistic model showed that whether novelty will be consciously accessed can be predicted by faster reaction times and increased amplitude of the P3 (table 1). This means that stimulus-driven phenomena (i.e., stimulus evaluation, indexed by the P3a, and overt decision-making processes, indexed by RT) account for increased odds of novelty being consciously accessed. Conversely, increased pupil size is associated with decreased probability of accessing sensory novelty. Such results illustrate the trade-off between bottom-up sensory processing and top-down mechanisms during goal-oriented behavior and highlight the role of NE-mediated arousal adaptation in weighing neural responses to novelty and uncertainty (Aston-Jones & Cohen, 2005; Joshi et al., 2016; Vazey et al., 2018; Zhao et al., 2019).

### Limitations

Our analyses are restricted to latencies associated with two classical ERP components, but the neural events underlying the processing of sensory novelty need not be limited to these time windows. This analytic decision was made so that new results could be related to and framed within well-grounded literature on conscious access to auditory novelty (Bekinschtein et al., 2009; Comerchero & Polich, 1999; Garrido et al., 2009; Näätänen et al., 2007, 2011; Polich, 1987, 2007). It could also be argued that a variable number of standard tones preceding each target could result in an anticipatory effect that may have confounded pupil results. Although such possibility cannot not be completely ruled out, control analyses showed no evidence that the number of preceding standard tones systematically modulated pupil responses across conditions (supplementary materials, figure s2). Another caveat is that we exclusively focused on the role of theta oscillations, as they have been previously implicated in novelty detection and related to the MMN and the P3. However, other studies have proposed a role of alpha oscillations in gating conscious perception and beta oscillations in prediction-error (Arnal et al., 2015; Chang et al., 2016; El Karoui et al., 2015; Haque et al., 2020). We think that looking into the role and relationship between other oscillatory bands and mechanisms during conscious access to novelty detection would be an insightful venue for future research. Finally, the staircase procedure used to identify discrimination thresholds could be improved, for instance, by using logarithmically spaced frequencies that are more faithful to human auditory perception or that better represent tonotopic representations in the cochlea.

### Conclusion

This study shows how the brain responds to suprathreshold stimuli that are consciously accessed as novel and to subthreshold novel stimuli that are subconsciously processed, but not consciously accessed. Behavioral results and replication of well-established ERP findings lend suggest that our experimental paradigm successfully teased apart conscious from subconscious processing of sensory novelty. Increased theta power reflects information maintenance about consciously accessed novel stimuli, which would be required for subsequent accurate decision-making. Conscious access to novelty is predicted by bottom-up markers of novel stimulus evaluation and overt decision-making processes. In contrast, subconscious processing of novelty is associated with increased phasic arousal adaptation due to heightened sensory uncertainty. These results highlight the dynamic relationship between bottom-up and top-down mechanisms and how the brain weights neural responses to novelty and uncertainty during perception and goal-directed behavior.

## Acknowledgments

We would like to acknowledge Dr. Rodrigo Henriquez-Ch, Dr. Gonzalo Boncompte, Dr. Vicente Medel, Dr. Marcos Domic, Dr. Pablo Billeke, Dr. Eugenio Rodriguez, Dr. Vladimir López and Dr. Brice Follet for their feedback and insightful comments and suggestions throughout the project.

## Author contributions

**Sergio Osorio:** conceptualization, methodology, formal analysis, validation, software, data curation, investigation, project administration, writing – original draft, writing – review & editing. **Martin Irani:** methodology, formal analysis, software, writing – review & editing. **Javiera Herrada:** validation, data curation, investigation, project administration, **Francisco Aboitiz:** conceptualization, resources, writing – review & editing, supervision, funding acquisition.

## Data and code availability statement

Access to data and scripts can be granted upon request.

