## Supplementary materials for "Neural responses to sensory novelty with and without conscious access"

**This PDF includes:**

Figures s1 and s2

Tables s1 and s2

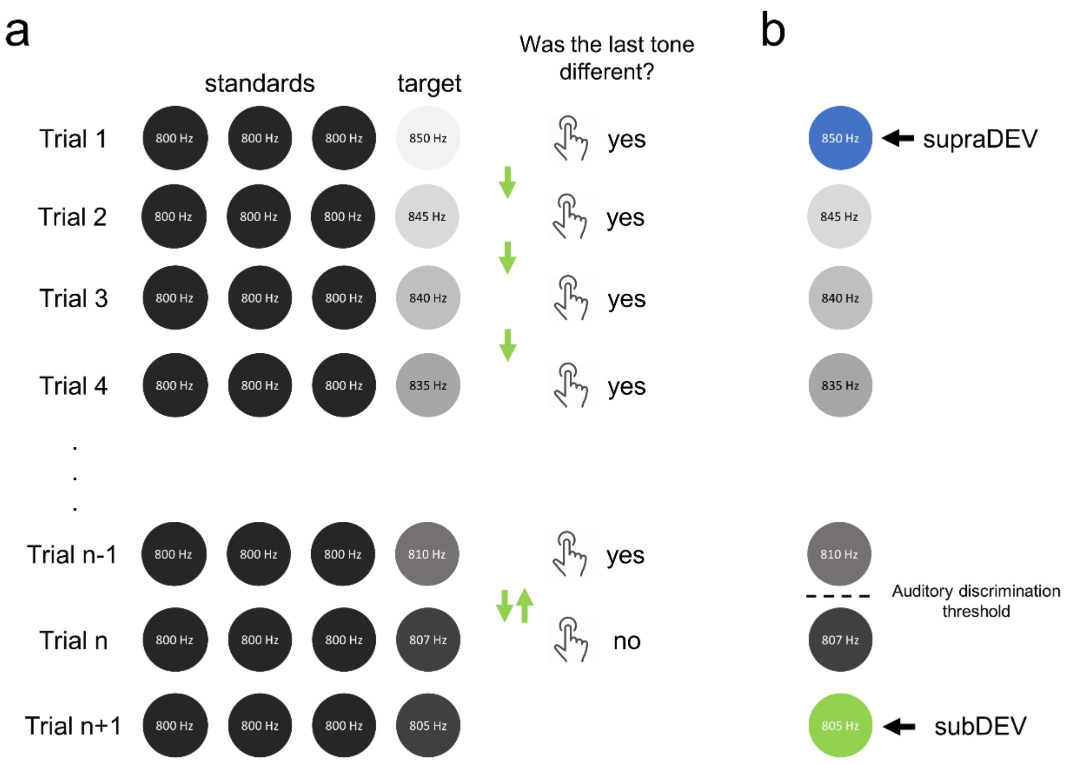


**Figure s1**. Staircase procedure. **a.** Sequences of standard tones (represented by black circles) with a base frequency of 800, 1000 or 1200 Hz, depending on the block, were presented to participants. In each trial, the number of standard tones before each target randomly varied between three and five tones. The first target was 50 Hz above the base frequency. Participants were asked whether the target was the same or different from the previous tones. Every time participants judged the target as different, the target was stepped down 5Hz in the next trial (green arrows, but see table s2). As the target approach participants discrimination threshold, they would eventually judge the target as same, even though it was not. At this point, the target was stepped up 5Hz in the next trial. At threshold, participants would enter a "same-different" loop (upward and downward green arrows). **b.** After five iterations of this loop, the discrimination threshold was defined as the imaginary boundary halfway between these two target frequencies, and the subDEV stimulus was set to be the second stimulus below that boundary. This process was carried out per subject, and per block.

| Block 1 | Block 2 | Block 3 |
|---|---|---|
| 850 | 1250 | 1050 |
| 845 | 1245 | 1045 |
| 840 | 1240 | 1040 |
| 835 | 1235 | 1035 |
| 830 | 1230 | 1030 |
| 825 | 1225 | 1025 |
| 820 | 1220 | 1020 |
| 815 | 1215 | 1015 |
| 810 | 1210 | 1010 |
| 807 | 1207 | 1007 |
| 805 | 1205* | 1005 |
| 804* | 1204 | 1004* |
| 803 | 1203 | 1003 |
| 802 | 1202 | 1002 |
| 801 | 1201 | 1001 |
| 800 | 1200 | 1000 |

**Table s1**. Stimuli sets for the staircase procedure. A different set of stimuli was used for each block. Stimuli were 5Hz apart from each other down to the base frequency plus 10 Hz. Because some people can show sharp hearing abilities, the staircase jumped from 10 to 7hz, and then decreased in steps of 1Hz. Asterisks represent the mode for subDEV targets across participants.

| Model | Formula | AIC | BIC |
|---|---|---|---|
| 1 | Conscious access ~1 + RT diff + pupil diff + mean MMN + mean P3 + mean power MMN + mean power MMN diff + mean power P3 + mean power P3 diff | 41.780 | 52.544 |
| 2 | Conscious access ~1 + RT diff + pupil diff + mean P3 + mean power MMN + mean power MMN diff + mean power P3 + mean power P3 diff | 38.693 | 48.852 |
| 3 | Conscious access ~1 + RT diff + pupil diff + mean P3 + mean power MMN + mean power MMN diff, mean power P3 | 35.861 | 45.239 |
| 4 | Conscious access ~1 + RT diff + pupil diff + mean P3 + mean power MMN diff + mean power P3 | 35.861 | 45.239 |
| 5 | Conscious access ~1 + RT diff + pupil diff + mean P3 + mean power MMN diff | 32.059 | 39.401 |
| 6 | Conscious access ~1 + RT diff + pupil diff + mean P3 | 30.920 | 37.031 |
| 7 | Conscious access ~1 + RT diff + pupil + mean P3 | 25.752 | 31.863 |
| 8 | Conscious access ~1 + RT + pupil + mean P3 | 35.280 | 41.391 |
| 9 | Conscious access ~1 + RT + pupil diff + mean P3 | 31.910 | 38.021 |
| 10 | Conscious access ~1 + RT diff + pupil | 44.788 | 49.541 |
| 11 | Conscious access ~1 + RT diff + mean P3 | 29.890 | 34.643 |
| 12 | Conscious access ~1 + pupil + mean p3 | 38.269 | 43.021 |
| 13 | Conscious access ~1 + RT diff | 49.412 | 52.688 |
| 14 | Conscious access ~1 + pupil | 63.086 | 66.362 |
| 15 | Conscious access ~1 + mean P3 | 38.255 | 41.531 |

**Table s2.** Model comparison. The formula for each model is presented along with their corresponding Akaike Information Criterion (AIC) and Bayesian Information Criterion (BIC).

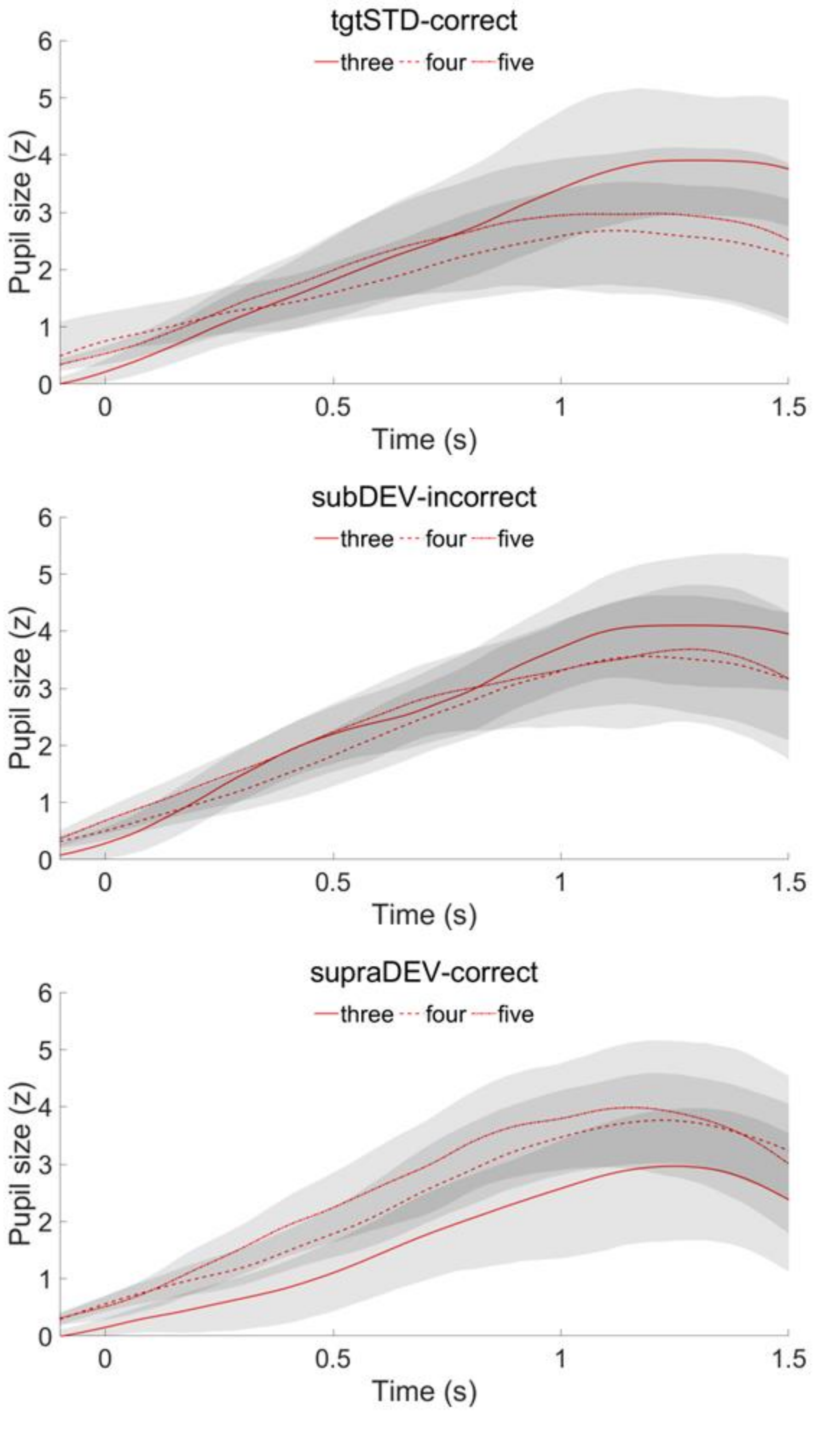


**Figure s2.** Pupil responses as a function of number of preceding standard tones for tgtSTD-correct, subDEV-incorrect, and supraDEV-correct trials. Gray shades represent the 95% C.I. of the mean.
